# Filament assembly confers the mechanical resilience of the IFI16 restrictosome

**DOI:** 10.64898/2026.10.30.763160

**Authors:** Christina M. Stallings, Archit Garg, Jungsan Sohn

## Abstract

Interferon-inducible protein 16 (IFI16) is a mammalian innate immune sensor composed of a filament-forming pyrin domain (PYD) followed by tandem HIN200 domains that bind nucleic acids. First recognized as a major regulator of inflammatory responses against cytosolic double-stranded (ds)DNA, IFI16 filaments have been implicated in several key mechanical processes involving both host and pathogens, including viral replication restriction, latency regulation, replication-stress responses, and DNA-damage responses. Moreover, IFI16 filaments are ultra-stable, a key trait believed to drive its autoantigenicity. Although recent studies reported that IFI16 undergoes liquid-liquid phase separation (LLPS) via disordered linkers and multivalent protein•dsDNA interactions, whether such a metastable entity provides a framework for the mechanical activities of IFI16 remains unknown. Here, we show that PYD-driven filament assembly confers the mechanical stability of IFI16 condensates by compacting DNA into ultra-stable tangles capable of withstanding forces sufficient to induce duplex deformation. By contrast, HIN200•dsDNA complexes are much more sensitive to ionic perturbation, DNA pretension, and mechanical interference. Together, we demonstrate that filament assembly allows IFI16 to act as a mechano-sensitive restrictor.

## Introduction

Assembling supra-molecular signaling platforms has emerged as a central theme in innate sensing of danger-and pathogen-associated molecular patterns (D/PAMPs) ^1–6^. Upstream sensors remain monomeric while dormant; however, upon directly binding D/PAMPs, these multi-domain receptors undergo homo-oligomerization, often resulting in either disc-shaped or filamentous signaling platforms ^1–6^. Such higher-order assemblies then recruit downstream signaling partners by multi-partite protein•protein interactions, triggering switch-like responses ^1–8^. Notably, recent studies indicated that these supra-structures undergo phase-transition, even operating as functional solids ^4,9^.

Interferon-inducible protein 16 (IFI16) is a member of absent-in-melanoma-2 (AIM2)-like receptors (ALRs), which are defined by the presence of a pyrin-domain (PYD) and one or two nucleic-acid-binding HIN200 domains separated by long unstructured linkers (100 and 50 a.a. for IFI16; HIN200: hematopoietic interferon inducible nuclear antigen with 200 a.a.; Fig. 1A) ^10–13^. IFI16 detects various dysregulated nucleic acids arising from pathogenic conditions, with its most prominent mechanism of action being assembling filaments on double-stranded (ds)DNA ^14–22^. For example, in resting cells, IFI16 is closely associated with the host chromatin in the nucleus ^19–22^ where filament assembly is prohibited as nucleosomes hamper its one-dimensional (1D)-diffusion assisted oligomerization on dsDNA ^16^. Upon the emergence of cytosolic dsDNA, such as those arising from damaged mitochondria, and viral/bacterial infection ^14,15,20,22–24^, IFI16 is exported from the nucleus and assembles filamentous signaling complexes on unchromatinized dsDNA. Similarly, upon the emergence of unchromatinized DNA in the nucleus caused by either DNA damage in the host chromatin or invasion by viral DNA (e.g. herpes simplex virus, HSV), IFI16 assembles filaments on such naked, contiguous dsDNA ^19,21,22,25,26^. IFI16 then partakes in eliciting and balancing various innate immune responses, including accentuating cGAS-STING-dependent interferon signaling pathways and counteracting the AIM2-dependent cell death pathway ^20–22,24,27–30^.

**Figure 1.**
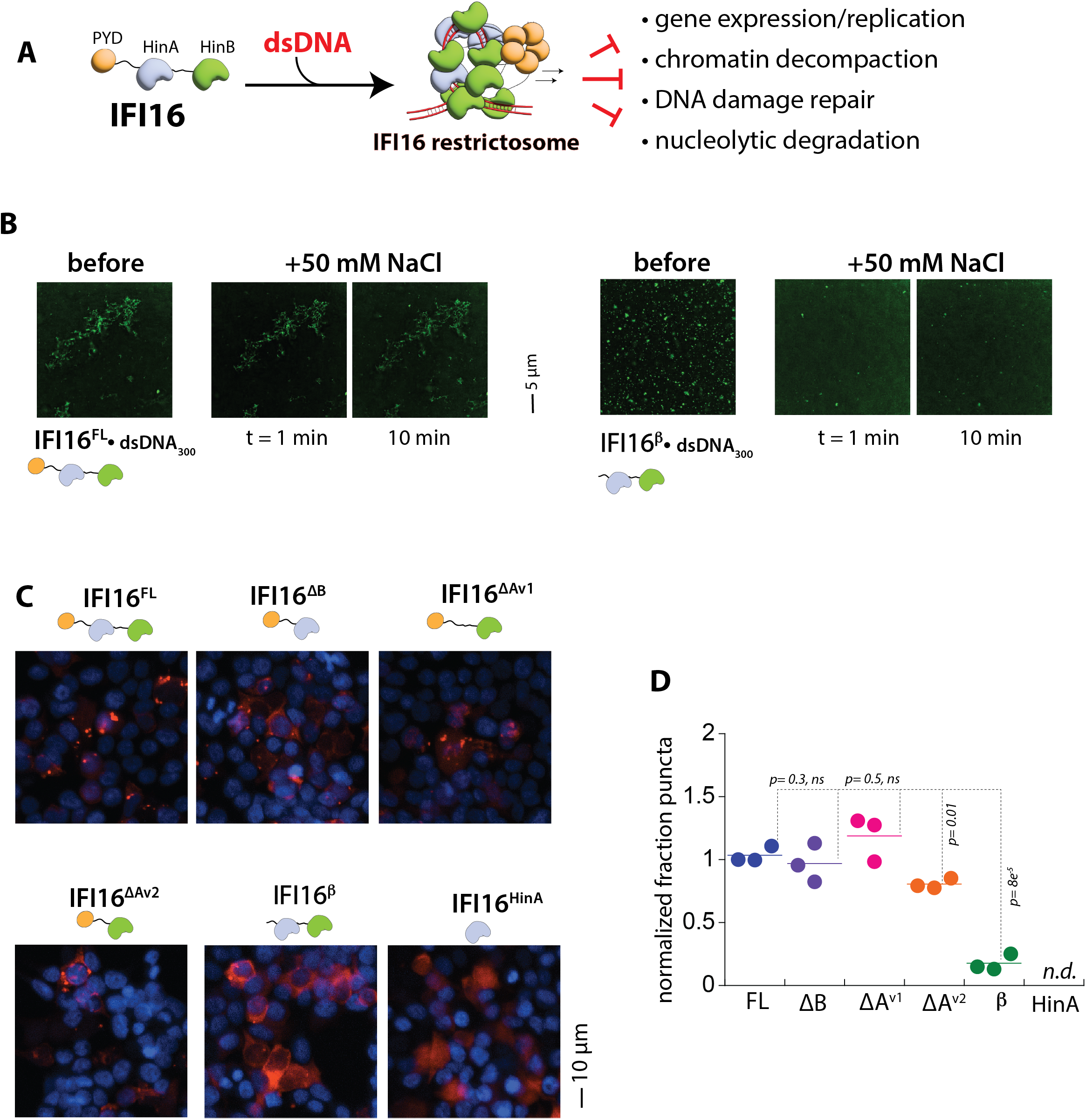
**A.** A cartoon describing the domain organization of IFI16 and its function as a restrictosome. **B.** Fluorescence images of IFI16^FL^ and IFI16β condensates formed on Alexa_488_-labeled 300-bp dsDNA before and after adding additional NaCl. **C.** Images showing C-terminally mCherry-tagged IFI16 variants expressed in HEK293T cells. ΔB: truncating after the first HIN200 domain. ΔA^v1^: deleting the first HIN200 domain while retaining the flanking linkers/IDRs. ΔA^v2^: replacing the first HIN200 domain with the second HIN200 domain then immediately truncating. HinA: the first HIN200 domain alone. Blue stains: DAPI. **D.** The fraction of puncta (condensates) formed per transfected cells relative to full-length IFI16 in HEK293T cells. ***ns***: not significant. ***n.d***.: not determined due to the lack of puncta formation.

In addition to its signaling function, a growing body of evidence suggests that IFI16 acts as a “restrictosome” (Fig. 1A). For example, IFI16 assembles into an ultra-stable filamentous complex on viral DNA, forming an immobile barrier to replication and transcriptional machineries ^22,25,31–34^. Through this mechanism, IFI16 restricts viral propagation, and also regulates the establishment and maintenance of latency (e.g., HSV, Epstein-Barr virus, and human papillomavirus) ^22,25,31–34^. Moreover, IFI16 filaments assembled on damaged host dsDNA restrict chromatin decompaction during DNA damage responses, consequently regulating tumor formation ^22,31,35,36^. Additionally, IFI16 filaments can shield host DNA by restricting nucleolytic degradation of stalled replication forks ^37^ and neutrophil extracellular traps (NETs) ^38^. However, the same ultrastability that enables IFI16 to form a robust restriction factor also predisposes it to elicit autoimmunity. Notably, long-lived IFI16 oligomers can persist as autoantigens in Sjögren syndrome and systemic lupus erythematosus, with the presence of anti-IFI16 antibodies directly correlating with increased disease severity ^15,38^.

Recent studies showed that IFI16 forms condensates on dsDNA via liquid-liquid phase separation (LLPS), thereby concentrating signaling components and accelerating immune activation ^39–41^. Yet, a sensor that merely condenses on DNA while remaining in the liquid phase faces many physical challenges to function as a robust restrictosome. For example, DNA, whether originating from the host or pathogens, is under constant mechanical tension ^42,43^: polymerases and replication forks exert forces on DNA templates, chromatin remodelers apply torque, and nucleosomes add curvature stress. A liquid condensate by nature is a highly dynamic and reversible entity ^44–46^, thus lacking resilience or durability required to compete against many other machineries targeting DNA. Currently, how IFI16 resolves this issue remains poorly understood. Here, we show that PYD filament assembly drives formation of a force-resistant, DNA-entangling supramolecular structure that physically immobilizes DNA and withstands mechanical interferences.

## Results

### Filament assembly bypasses the multivalency requirement for condensate formation

Recent studies have shown that the HIN200 domains and intrinsically disordered region (IDR) of IFI16 are sufficient to invoke LLPS with dsDNA, providing a dense metastable intermediate state for filament nucleation and antiviral activities ^40,41^. However, we previously found that the HIN200 domains bind dsDNA weakly and continuously 1D-diffuse on the duplex until cooperative filament assembly by multiple proximal protomers stabilizes binding to form immobile clusters ^16,17^. Notably, 1D-diffusion retains many IFI16 protomers in close proximity without repeated 3D searches, thereby raising their effective local concentration and the rate of specific PYD•PYD interactions required to nucleate and elongate filaments ^16–18,47,48^. LLPS therefore is not required under this mechanism.

To interrogate how filament assembly and LLPS are related, we tracked the condensate formation by full-length human IFI16^(FL)^ and its naturally occurring PYD-less isoform termed IFI16β ^49^. Here, both recombinant proteins produced large condensates with Alexa_488_-labeled 300-bp dsDNA (Fig. 1B). However, the full-length protein produced filamentous clusters, while those formed by the PYD-less variant appeared more droplet-like. Notably, introducing additional 50 mM NaCl did not affect the filamentous condensates formed by IFI16^FL^, but nearly abolished IFI16β•dsDNA condensates within one minute (Fig. 1B). These results indicate that IFI16β condensates are formed largely via multivalent electrostatic interactions between HIN200 domains and dsDNA, while the highly ordered PYD filament assembly ^48^ confers chemical resistance.

We next tracked the oligomerization of various mCherry-tagged IFI16 variants when transfected into HEK293T cells. Because wild-type IFI16 diffusively localizes in the nucleus when transfected into these cells ^20,50^, we incorporated a mutation mimicking acetylation to induce cytosolic export (K128Q) ^20^. Consistent with our previous observations ^48^, IFI16^FL^ and PYD-containing variants formed puncta, indicative of higher-order assemblies/condensates formed with transfected DNA (Fig. 1C). Although IFI16^ΔAv^^2^ produced fewer puncta, it was only ∼19% fewer than those produced by the full-length protein (Fig. 1C-D). However, the number of puncta produced by IFI16β was ∼82% fewer than those seen from IFI16^FL^, and IFI16^HinA^ completely failed to produce any puncta (Fig. 1C-D). Our observations indicate that PYD-driven filament assembly results in a phase transition, providing chemical resistance necessary to form condensates in cells.

### Filament assembly entangles dsDNA and confers the mechanical stability for the IFI16 restrictosome

While our ensemble experiments reveal the morphological and compositional requirements for condensate formation, they cannot report on the physical forces that hold these assemblies together, any potential mechanical work IFI16 might perform on dsDNA, or how individual complexes respond to forces imposed on dsDNA. We thus employed an optical tweezer (Lumicks C-Trap) to further probe the mechanical basis for the condensate formation by IFI16.

We first asked whether IFI16 binding generates force (i.e., performs mechanical work) on λdsDNA tethered at a fixed distance between two optical traps (10 µm, no tension). Incubating trapped λdsDNA in a microfluidics chamber containing various concentrations of IFI16^FL^ generated up to ∼3 pN of force on λdsDNA within 200 sec (Fig. 2A-C; 0.05, 0.1, 0.2 and 1 µM; K_D_ for 300-bp < 10 nM ^16,17^). Considering the much faster dsDNA binding kinetics of IFI16 (t_1/2_ < 10 sec) ^16^ and the concentration independent increase in force beyond 100 nM, these results likely reflect secondary mechanical processes, not simply protein loading onto λdsDNA. Supporting this notion, comparing the force tracking with the kymograph of Cy3-labeled IFI16^FL^ showed that the rise in force coincides with the accumulation of IFI16 on λdsDNA followed by formation of immobile clusters (Fig. 2C). Our observations are also consistent with our previous findings using flow-stretched λdsDNA in which individual IFI16 molecules randomly diffuse along λdsDNA and form immobile oligomers once 7 to 8 protomers come into proximity ^16^. Moreover, these results reveal that the oligomerization of IFI16 on dsDNA is a force-dependent, mechanical process.

**Figure 2.**
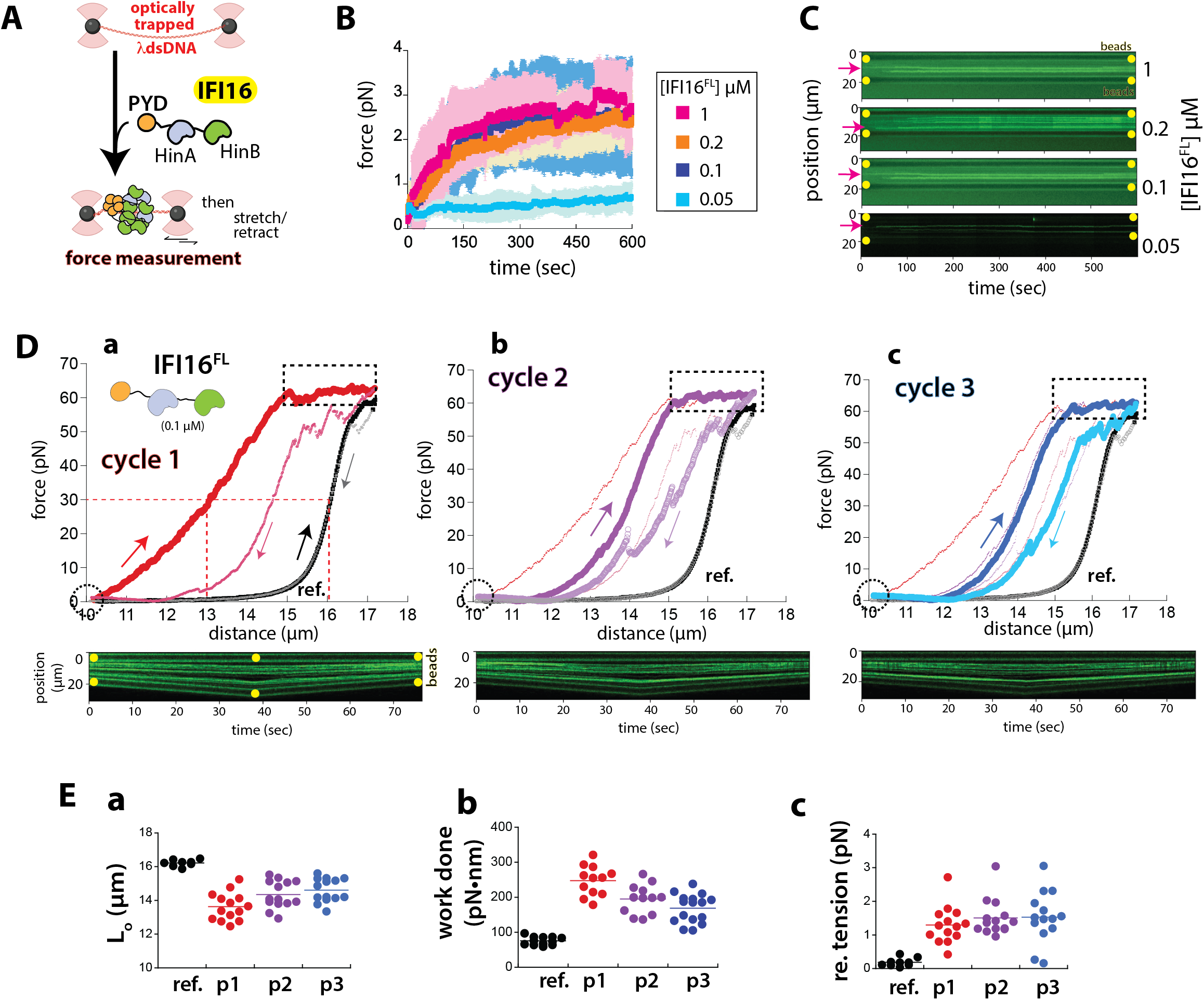
**A.** A cartoon describing passive mode and force extension experiments using λdsDNA tethered by the dual trap in the Lumicks C-Trap. **B.** Average force readings over time upon introducing indicated concentrations of IFI16^FL^ to slack λdsDNA. *n* ≥ 5, ± standard deviations. **C.** Representative confocal scans (kymographs) of λdsDNA upon adding indicated concentrations of IFI16^FL^. Bead positions are indicated with yellow dots. Magenta arrows point at example traces indicative of IFI16^FL^ accumulation on λdsDNA. **D.** Representative extension and retraction curves of IFI16^FL^-bound or naked λdsDNA. Diagonally upward arrows and filled data points (thicker colors) indicate extension, while downward arrows and open data points (lighter colors) indicate retraction. For **b** and **c**, data from Cycle 1 and 1/2 are shown as thinner lines, respectively. ref: λdsDNA alone. **E.** Dot plots showing the contour length (L_0_), work done, and remaining tension on λdsDNA after pull (p) 1, 2, and 3.

At the end of passive-mode experiments (10 min incubation), we moved each IFI16-bound λdsDNA to the buffer only channel and stretched it three consecutive times (as illustrated in Fig. 2A). We also simultaneously tracked the changes in force and visualized bound complexes via the confocal function of the C-Trap (i.e., force extension (FE) experiments; Fig. 2D). Here, stretching IFI16-bound λdsDNA immediately required more force than the naked counterpart (Fig. 2D-a), which is consistent with the behaviors expected from entangling/bridging tethered dsDNA by bound proteins ^51–55^. Indeed, comparing the pulled distance at 30 pN ^56^ indicated that the contour length of λdsDNA in complex with IFI16 was ∼ 2.5 µm shorter than the unbound counterpart (red dashed lines in Fig. 2D-a and ref. vs. p1 in Fig. 2E-a), which also correlated with the significantly more work required for pulling the tethered λdsDNA (ref. vs. p1 in Fig. 2E-b). Notably, λdsDNA remained largely contracted even when it was experiencing force-induced deformation ^51–55^ (Fig. 2D-a, dotted box). Moreover, the retraction curve (open circles in Fig. 2D-a), albeit considerably right-shifted than the extension curve, still largely deviated from that of naked λdsDNA, again indicating that λdsDNA is still contracted. Visually tracking IFI16 during extension and retraction via the confocal function showed that although some bound proteins are indeed lost during the pull, many remained (kymographs shown on the bottom of Fig. 2D-a). Stretching the same bound complex two more times showed that ∼0.8 µm of λdsDNA had been liberated by the initial cycle (Fig. 2D-b/c and p1 vs. p2/p3 in Fig. 2E-a). However, λdsDNA was still tangled and contracted by bound IFI16^FL^ clusters (Fig. 2D-b/c and ref. vs. p2/p3 in Fig. 2E-a). Notably, the second and third cycles produced essentially identical FE profiles, as λdsDNA remained contracted by bound IFI16 clusters even after repeatedly experiencing duplex deformation (Fig. 2D-b/c and p2 vs. p3 in Fig. 2E-a/b). Further consistent with persistent dsDNA entanglement, the force reading on λdsDNA remained significantly higher after being completely retracted (dotted circles in Fig. 2D and Fig. 2E-c). Together, our experiments reveal that IFI16 entangles dsDNA to assemble into highly force-resistant clusters.

To determine whether the observed force exertion and resistance arise from filament assembly, we examined the mechanical properties of λdsDNA upon binding the truncated variants we used in Fig. 1. IFI16 constructs retaining the PYD showed force profiles comparable to the full-length protein. For instance, in the passive mode, they generated up to 4 pN of force upon binding to slack λdsDNA when present at saturating concentrations (Fig. 3A). Subsequently stretching the bound complexes also immediately required greater force than the unbound counterpart, indicating that λdsDNA is contracted and stiffened into tangles (Fig. 3B). However, unlike IFI16^FL^, force-extension curves from these constructs showed conspicuous dips during the first pull (dotted boxes in Fig. 3B), suggesting that loops formed by constructs lacking either HIN domain are larger than those generated by the full-length protein. These observations also suggest that the resulting complexes are not as force resistant as those formed by the full-length due to the missing HIN domain. Nevertheless, subsequent cycles showed that many IFI16 clusters are still bound and withstood duplex deformation force (magenta arrows in Fig. 3B); λdsDNA also remained contracted and tensed (dotted circles in Fig. 3B). Notably, the amount of contracted dsDNA and work necessary to pull bound complexes were comparable to those seen from IFI16^FL^ in both the initial and last cycles (Fig. 3C-a/b; 3C-c/d), and λdsDNA also remained similarly tensed after three cycles of extension and retraction (Fig. 3C-e). Comparing average first pulls with one another also showed that these PYD-containing constructs are as competent as IFI16^FL^ in contracting dsDNA into force-resistant tangles. These results consistently suggest that PYD-driven filament assembly is critical for mechanical force resistance, compaction, and dsDNA entanglement.

**Figure 3.**
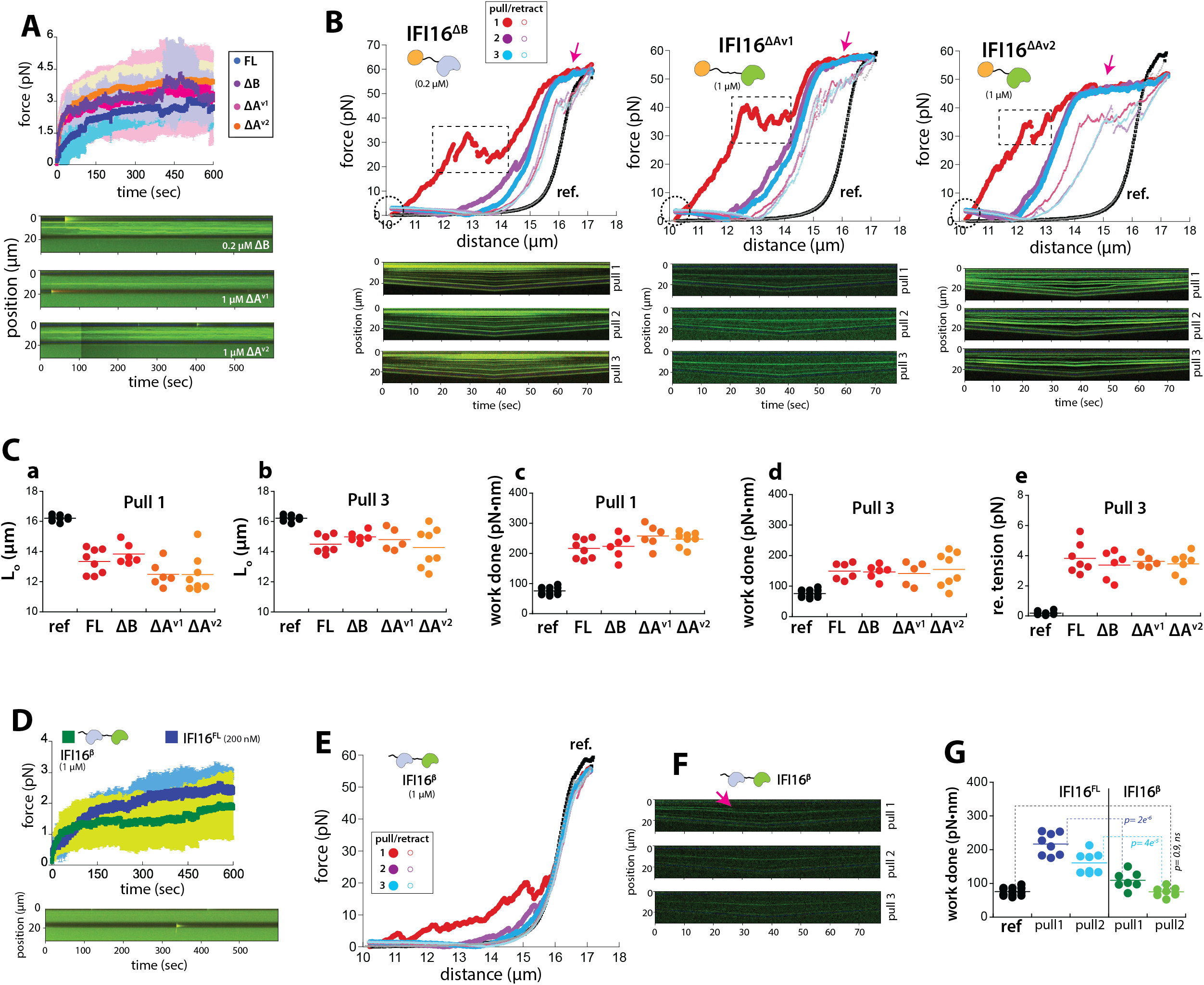
**A. Top**: Average force readings over time upon introducing 1 µM of indicated IFI16 constructs to slack λdsDNA. *n* ≥ 5, ± standard deviations, ref.: λdsDNA alone. **Bottom**: representative kymographs showing λdsDNA upon introducing each IFI16 construct. **B.** Representative extension and retraction curves of unbound λdsDNA or bound with each IFI16 construct. Filled data points (thicker colors) indicate extension, while open data points (thinner/smaller colors) indicate retraction. **Bottom**: corresponding kymographs taken during each cycle. Dotted boxes indicate conspicuous dips during the first pull. **C.** Dot plots showing the contour length (L_0_), work done, and remaining tension on λdsDNA when bound to different IFI16 constructs. Pull cycle is also indicated. **D. Top**: Average force readings over time upon introducing IFI16^FL^ or IFI16β to slack λdsDNA. *n* ≥ 5, ± standard deviations. **Bottom**: a representative kymograph showing λdsDNA upon introducing IFI16β. **E.** Representative extension and retraction curves of unbound λdsDNA or bound with the IFI16β. Filled data points (thicker colors) indicate extension, while open data points (lighter/smaller colors) indicate retraction. ref.: λdsDNA alone. **F.** Corresponding kymographs taken during each pull cycle shown in **E**. The magenta arrow indicates the loss of fluorescence signal (i.e. loos of bound IFI16β) while pulling bound λdsDNA. **G.** Dot plots showing the calculated work done while pulling λdsDNA bound to either IFI16^FL^ or IFI16β.

Although binding of IFI16β to slack λdsDNA also generated force, the magnitude was limited to less than 2 pN even at saturating/condensate-forming concentrations (1 µM, as seen in Fig. 1B; Fig. 3A vs. 3D). Pulling the bound complexes showed a modest rise in force accompanied by multiple dips and rises indicative of release of loops and protrusions ^51–55^ (Fig. 3E). Notably, these release events started to occur well below 10 pN (Fig. 3E), which is in contrast to the full-length protein (none in Fig. 2D) and the PYD-retaining constructs that lack either HIN200 domain (>20 pN; Fig. 3B). Moreover, virtually all topological changes on dsDNA made by the PYD-less construct were undone by the first pulling cycle (Fig. 3E), which was accompanied by confocal images showing proteins being dislodged during extension (e.g. magenta arrows in Fig. 3F). Additionally, the retraction profile became essentially identical to naked λdsDNA, and subsequent pull-and-retract cycles were largely indistinguishable from unperturbed λdsDNA (Fig. 3E). Our observations are expected from a complex held together primarily through electrostatic HIN200-mediated contacts ^16–18^. These individually weak and non-cooperative interactions are then irreversibly disrupted at modest forces, as the HIN200 domains continuously diffuse along the 1D scaffold ^16^. Indeed, substantially less mechanical work was necessary to stretch λdsDNA bound to IFI16β compared to those in complex with PYD-retaining constructs, and the amount of work required during the second pull was essentially indistinguishable from pulling the naked λdsDNA (e.g., Fig. 3G). Collectively, our experiments demonstrate that filament assembly by PYDs provides the foundation for assembling a highly force-resistant complex with dsDNA.

### Filament assembly is essential for IFI16 to overcome dsDNA pretension

dsDNA *in vivo*, whether from hosts or pathogens, is under constant tension ^42,43^. Given the mechanical resilience of IFI16 oligomers on dsDNA, we next asked whether the assembly can overcome the pre-existing tension on dsDNA. To test this idea, we monitored whether IFI16 can contract λdsDNA held at different tensions by the optical trap (Fig. 4A top). In these force clamp (FC) experiments, 100 nM IFI16 rapidly and completely contracted λdsDNA held at 1, 2, and 3 pN (Fig. 4A). However, IFI16 only marginally contracted λdsDNA held at 4 pN (∼2 µm), albeit binding readily (Fig. 4A). The variable intensity of IFI16 in the kymographs suggests that some oligomers are still formed on dsDNA held at higher tension (Fig. 4A); however, these locally formed oligomers are apparently unable to bridge longer distances to contract dsDNA against the pre-existing tension.

**Figure 4.**
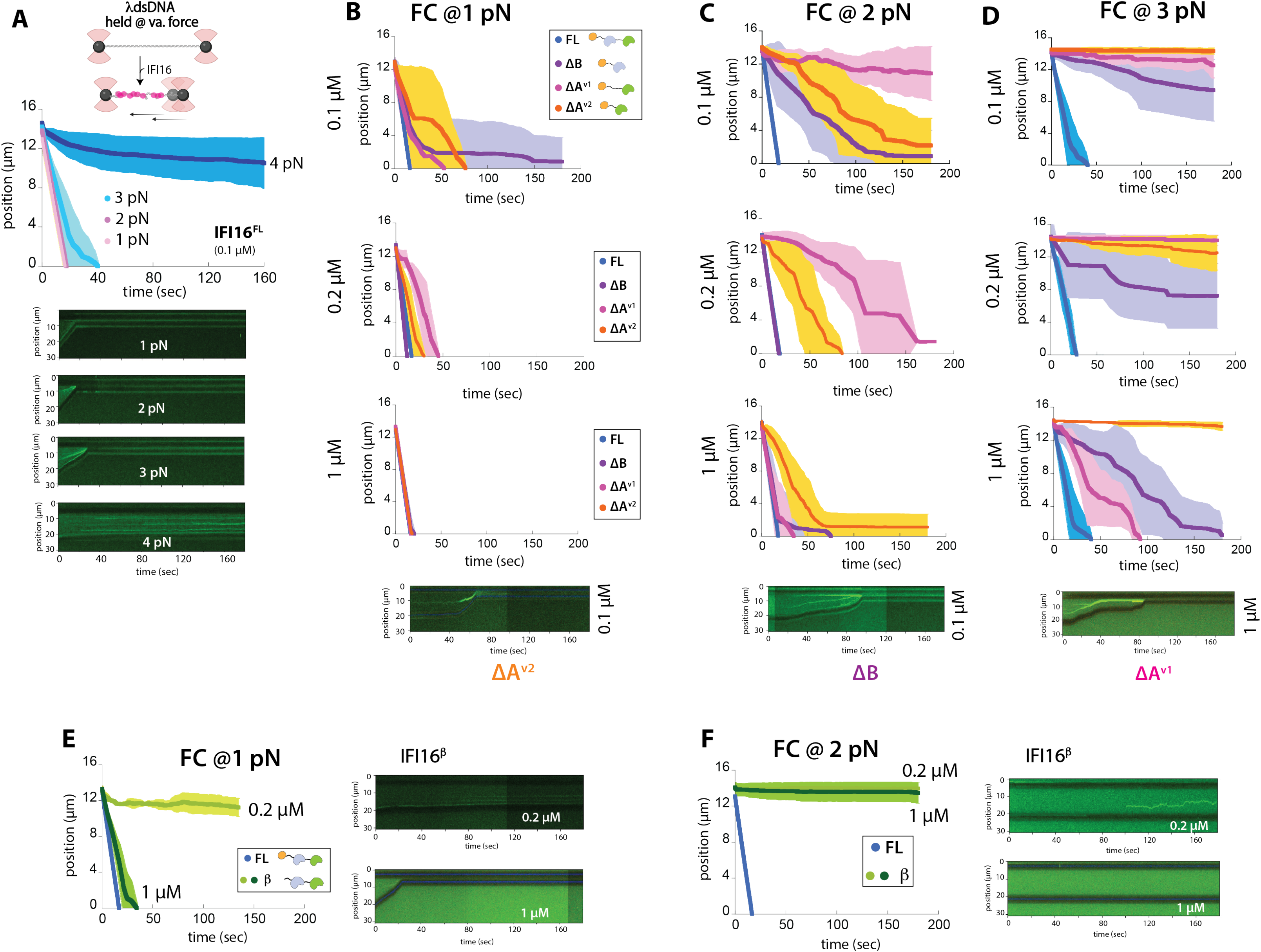
**A. Top**: a cartoon describing force clamp experiments using the Lumicks C-Trap. **Middle**: average position/distance between two traps (beads) over time upon introducing IFI16^FL^ to tethered λdsDNA held at indicated tension. *n* ≥ 5, ± standard deviations. **Bottom**: representative kymographs from λdsDNA held at indicated tension upon introducing IFI16^FL^. **B.** Average position/distance between two traps (beads) over time upon introducing each IFI16 construct at indicated concentration to tethered λdsDNA held at 1 pN. *n* ≥ 5, ± standard deviations. A representative kymograph of IFI16^ΔAv2^ is shown on the bottom. **C.** Average position/distance between two traps (beads) over time upon introducing each IFI16 construct at indicated concentration to tethered λdsDNA held at 2 pN. *n* ≥ 5, ± standard deviations. A representative kymograph of IFI16^ΔB^ is shown on the bottom. **D.** Average position/distance between two traps (beads) over time upon introducing each IFI16 construct at indicated concentration to tethered λdsDNA held at 3 pN. *n* ≥ 5, ± standard deviations. A representative kymograph of IFI16^ΔAv1^ is shown on the bottom. **E.** Average position/distance between two traps (beads) over time upon introducing IFI16^FL^ or IFI16β to tethered λdsDNA held at 1 pN. *n* ≥ 5, ± standard deviations. Representative kymographs of IFI16β bound λdsDNA are shown on the right. **F.** Average position/distance between two traps (beads) over time upon introducing IFI16^FL^ or IFI16β to λdsDNA held at 2 pN. *n* ≥ 5, ± standard deviations. Representative kymographs of IFI16β bound λdsDNA are shown on the right.

Increasing the concentrations of IFI16^FL^ to 0.2 and 1 µM essentially showed the same trend as the results from 0.1 µM (Fig. 4B-D). Using these conditions, we then examined PYD-retaining constructs (Fig. 4B-D). At 1 pN, except at 0.1 µM, IFI16^ΔB^ contracted λdsDNA as readily as the full-length protein (Fig. 4B). The two IFI16^ΔA^ variants showed slightly slower kinetics at lower concentrations, but compacted λdsDNA held at 1 pN as well as IFI16^FL^ at the saturating concentration (1 µM; Fig. 4B). At 2 pN, the compaction by IFI16^ΔB^ was further slowed for 0.1 µM (t_1/2_ ∼ 65 sec vs. ∼ 5 sec for IFI16^FL^), but higher concentrations resulted in full-length-like compaction activities (0.2 and 1 µM in Fig. 4C). The IFI16^ΔA^ variants compacted dsDNA more slowly still: 0.1 µM of ΔAv1 failed to compact dsDNA held at 2 pN entirely, while higher concentrations progressively restored activity (Fig. 4C). Similarly, the compaction activity of ΔAv2 also increased with the higher concentrations (Fig. 4C). At 3 pN, the gap between ΔB and the full-length protein widened further, with the two lower concentrations of ΔB failing to fully contract λdsDNA (Fig. 4D). Moreover, at lower concentrations, both IFI16^ΔA^ variants failed to contract λdsDNA despite binding (Fig. 4D). At 1 µM, ΔAv1 compacted λdsDNA while ΔAv2 still failed (Fig. 4D), suggesting that the nonspecific interactions from the additional IDR assisted in oligomerization once fully bound. Nevertheless, IFI16^FL^ contracted λdsDNA held at 3 pN most readily (Fig. 4D).

Unlike the PYD-retaining constructs, although 0.2 µM of IFI16β bound λdsDNA held at 1 pN, it failed to contract the scaffold; increasing the concentration to 1 µM allowed it to contract λdsDNA (Fig. 4E). Nevertheless, the PYD-less construct failed to compact λdsDNA when the tension was raised to 2 pN (Fig. 4F). Notably, kymographs suggest that IFI16β fails to form any stable complexes/clusters even with a small increase in tension (Fig. 4F). Collectively, our observations suggest that PYD filament assembly is essential for IFI16 to overcome any significant pretension on dsDNA, with the tandem HIN200 domains acting cooperatively to further facilitate these processes.

## Discussion

IFI16 is a versatile innate immune sensor that partakes in a wide range of host responses, which include type-I interferon (IFN-I) activation, pathogen replication restriction, and viral latency regulation ^14–22^. Moreover, IFI16 regulates several key cellular processes such as cellular senescence, DNA damage responses, and tumor formation ^35,57,58^. At the crux of all these different facets of IFI16 is filament assembly in which its HIN200 domains use a contiguous stretch of unchromatinized dsDNA as a 1D-diffusion scaffold to locate one another to promote the oligomerization of PYDs ^16–18^. Recent studies proposed that IFI16 forms liquid-like condensates with dsDNA via IDR and multivalent protein•dsDNA interactions, which plays a key role in its signaling function ^40,41^. However, LLPS condensates by nature are dynamic and metastable entities, thus unlikely to provide the stability required of a restriction factor. Here, our experiments reveal that filament assembly provides resistance and stability necessary for IFI16 to withstand environmental and mechanical perturbations.

### LLPS vs. Filament formation and phase-transition

It is increasingly appreciated that condensate formation is integral to many biological processes ^44,46,59,60^. Classically, LLPS was considered essential for generating functional biomolecular condensates, while gel-or solid-like phase-transitioned states represented disease states (e.g., amyloid-β solids ^44,46,59,60^). However, it is now recognized that much denser condensates can represent functional/active states ^44,46,59,60^. For example, cGAS forms gel-like condensates with cognate dsDNA substrate ^61,62^, while undergoing LLPS with other nucleic acids that do not activate the enzyme (e.g., dsRNA and single-stranded DNA/RNA) ^61,62^. Similarly, inflammasome receptors form gel-like condensates upon activation that further transition into solid-like functional aggregates ^4,9^. Here, we show that the filament assembly of IFI16 results in condensates that are both chemically and mechanically resilient. In fact, our experiments with the PYD-less construct suggest that filament assembly is critical for forming condensates in cells (Fig. 1C). It is noteworthy that IFI16 filament assembly occurs via 1D-diffusion on dsDNA ^16^, which does not necessitate LLPS. Thus, our results here suggest that establishing a percolative network ^59^ via filament assembly, but not segregative LLPS via transient HIN•dsDNA interactions ^44,46,59^, is the main driver for IFI16 condensates to attain the mechanical resilience necessary to act as restrictosomes.

### The mechanical underpinning of the IFI16 restrictosome

The experiments described here reveal that IFI16 is not merely a pattern recognition receptor that signals through downstream effectors, but a mechanical restrictor that physically subdues foreign and damaged DNA through force-resistant filament assembly. For instance, HIN200-mediated contacts impose modest topological changes on dsDNA; however, loops and protrusions formed by IFI16β are released sequentially and almost completely at forces well within the range of transcription and chromatin remodeling machineries (5–20 pN) ^63^ and are irreversibly disrupted within a single pulling cycle (Fig. 3E-F). By contrast, PYD filament assembly produces a completely different mechanical state: dsDNA is compacted by ∼2.5 µm, stiffened into force-resistant tangles, and remains contracted even after repeated cycles of duplex-deforming extension (Figs. 2-3).

It is also noteworthy that filament assembly is essential for IFI16 to compact dsDNA against physiological pretension ^42,43,56^, as IFI16 was able to overcome up to 4 pN of dsDNA pretension at least partially (Fig. 4A). The concentration dependence of compaction across truncated variants further reveals that the tandem HIN200 domains act cooperatively to reduce the force threshold for filament nucleation (Fig. 4B-D). Our data also suggest that the tighter binding affinity of IFI16^HinA^ confers a measurable kinetic advantage over HinB-only constructs (ΔB vs. ΔA variants; Fig. 4B-D), and the IDR in ΔAv1 provides additional nonspecific interactions that partially compensate for reduced HIN domain multivalency at higher pretension under saturating concentrations (Fig. 4D). These observations are consistent with a mechanistic model in which HIN200 domains bring IFI16 protomers in proximity on dsDNA, but PYD•PYD interactions provide the free energy necessary to overcome pretension and drive compaction.

We found here that IFI16^FL^ complexes can repeatedly resist forces that induce the deformation of the B-form DNA duplex (∼65 pN ^64,65^, Fig. 2). This is remarkable because such a force regime largely exceeds those seen in RNA polymerase stalling (∼30 pN) ^65^, nucleosome unwrapping (up to ∼40 pN) ^66^, and chromatin remodeling (∼12 pN) ^63^. Thus, viral polymerases or even helicases would not be able to mechanically displace the IFI16 restrictosome from episomal DNA. Likewise, for DNA damage responses, the ability of IFI16 to compact and immobilize damaged dsDNA under pretension suggests a direct physical mechanism for regulating chromatin decompaction at damage sites. For autoimmunity, the same mechanical stability that makes IFI16 an effective restriction factor explains the persistence of IFI16 oligomers as autoantigens in Sjögren syndrome and systemic lupus erythematosus ^15,38^, as such an irreversibly stable complex will not be cleared by routine turnover mechanisms that disassemble less stable complexes. On the other hand, our results show that dsDNA pretension can suppress the bridging of local clusters over longer distances (4 pN in Fig. 4A). Thus, it is tempting to speculate that dsDNA tension arising from various essential processes could physically suppress its spurious activation (e.g., cell division, replication, and gene expression).

Collectively, our findings establish that PYD-driven filament assembly confers the mechanical resilience and functional specificity of the IFI16 restrictosome, which would also underpin its exceptional persistence implicated in driving autoimmune disorders. This mechanical framework thus offers new therapeutic entry points for selectively destabilizing pathogenic IFI16 filaments in autoimmune disease without compromising its antiviral and regulatory roles.

## Acknowledgements

We thank Drs. Brendan Antiochos, Scott Bailey, Taekjip Ha, and Shuai Wu for discussion. This work was supported by NIH R21 AR085266, NIH R35 GM145363, NSF MCB 2501395, Johns Hopkins Synergy Award, and Jerome L. Greene Foundation Scholars Award to J.S.

## Author Contributions

C.M.S and J.S. conceptualized the project. C.M.S. and A.G. performed all experiments, analyzed data, and wrote the paper. J.S. supervised the overall project, analyzed data, and wrote the paper.

## Materials and Methods

All experiments were performed at least three times (biological replicates), and *p* values were calculated using the two-tailed *t-test* assuming unequal variances (Microsoft Excel). All error bars were obtained from calculating standard deviations (Microsoft Excel). Graphs were made using KaleidaGraph (Synergy) and Figures were organized using Adobe Illustrator.

### Recombinant proteins

Human IFI16^FL^ (residues 1-729) and other constructs containing PYDs were cloned into the pET21b vector with a C-terminal His_6_-tag. The IFI16β isoform (residues 159-729) was cloned into the modified pET28b vector with an N-terminal His_6_-SUMO tag, which was removed by the Ulp1 protease before all experiments. All plasmids were transformed into *E. coli* BL21-Rosetta 2^DE3^ cells and purified via Ni^2+^-NTA, cation exchange, and size exclusion chromatography (storage buffer: 40 mM HEPES-NaOH at pH 7.4, 400 mM NaCl, 1 mM TCEP, 1 mM EDTA and 10 % glycerol). Proteins were then concentrated and stored at -80°C. Each protein was labeled by the Cy3 fluorophore using maleimide chemistry (ThermoFisher). See also ref. ^16,17,67^.

### Condensate assays

Cellvis-384 Glass Bottom plates were cleaned with a 5% Hellmanex Solution, rinsed with deionized water (DI), and subsequently etched with 1 M KOH. The plates were then thoroughly rinsed with DI water, dried overnight, and sealed with aluminum foil. Prior to use, wells were unsealed and rinsed with reaction buffer (40 mM HEPES-NaOH at pH 7.4, 160 mM KCl, 1 mM TCEP, 1 mM EDTA and 5% glycerol). Unlabeled IFI16 proteins (1 µM) were mixed with 300-bp dsDNA (1 μM; binding sites normalized to 15-bp; 5% Alexa Fluor 488-labeled) in a final volume of 25 μl. Images were acquired every 5 min using a Cytation 5 imaging plate reader (Agilent). Images were analyzed for condensate formation using the BioTek Gen5 software (Agilent).

### Cell culture and imaging

Each IFI16 construct was cloned into a pCMV6 vector containing a C-terminal mCherry tag. Indicated plasmids were transfected into HEK293T cells at ∼70% confluency (12 well plates) using lipofectamine 3000 (Invitrogen). After 18 hours, cells were washed with 1x PBS, fixed with 4% paraformaldehyde followed by staining with Hoechst reagent (Invitrogen) and mounted on glass slides using ProLong Gold antifade reagent (Invitrogen). Imaging was performed using the Cytation 5 plate reader equipped with a fluorescent microscope (Agilent) and analyzed via Gen5 software (Agilent). The ratio of amount of IFI16 oligomers/cells were calculated for each construct and normalized to those seen from the full-length protein. At least 3 biological replicates were conducted.

### Single-molecule experiments

All single-molecule experiments were conducted using the C-Trap (Lumicks). The C2 microfluidics system was passivated using the following steps: 0.1% BSA in PBS flowed at 0.4 bar pressure for 30 min, followed by flowing PBS at 0.4 bar pressure for 10 min to rinse, followed by 0.5% Pluronics F-127 in PBS at 0.4 bar for 30 min, and finished with a final rinse of PBS at 0.4 bar for 30 min. 4.38 µm streptavidin coated polystyrene beads (Spherotech cat no. SVP-40-5) were captured in each trap and used to tether biotinylated λdsDNA (Lumicks). Experimental buffer contained 40 mM HEPES-NaOH at pH 7.4, 160 mM KCl, 1 mM TCEP, 1 mM EDTA, and 5% glycerol. Cy3-labeled proteins were mixed with their unlabeled counterparts with a 1:3 ratio.

In force extension (FE) experiments, tethered λdsDNA was first stretched by moving one of the captured beads at a constant pre-adjusted pulling speed of 0.2 µm/sec in the buffer only channel and the resulting force was recorded. The λdsDNA was then moved to a channel containing each IFI16 construct and incubated for 10 min where force reading was recorded (i.e., the passive mode). Subsequently, the λdsDNA was moved to the buffer only channel and stretched using a constant pulling speed of 0.2 µm/sec then the resulting force was recorded. In force clamp (FC) experiments, tethered λdsDNA was moved to the protein chamber and incubated for 3 minutes with the force clamp engaged at indicated force then data was recorded. We estimated the contour length of each IFI16•λdsDNA complex using the pulled distance at 30 pN ^56^. The work performed during each pull was calculated by integrating the area under each FE curve.

## Reference

1. Kagan, J.C., Magupalli, V.G. & Wu, H. SMOCs: supramolecular organizing centres that control innate immunity. Nat Rev Immunol 14, 821–6 (2014).

2. Kobe, B. et al. Signalling by co-operative higher-order assembly formation: linking evidence at molecular and cellular levels. Biochem J 482, 275–294 (2025).

3. Nanson, J.D., Kobe, B. & Ve, T. Death, TIR, and RHIM: Self-assembling domains involved in innate immunity and cell-death signaling. J Leukoc Biol 105, 363–375 (2019).

4. Huoh, Y.S. & Hur, S. Death domain fold proteins in immune signaling and transcriptional regulation. FEBS J 289, 4082–4097 (2022).

5. Sohn, J. & Hur, S. Filament assemblies in foreign nucleic acid sensors. Curr Opin Struct Biol 37, 134–44 (2016).

6. Lu, A. & Wu, H. Structural mechanisms of inflammasome assembly. FEBS J 282, 435–44 (2015).

7. Matyszewski, M., Morrone, S.R. & Sohn, J. Digital signaling network drives the assembly of the AIM2-ASC inflammasome. Proc Natl Acad Sci U S A 115, E1963–E1972 (2018).

8. Matyszewski, M. et al. Distinct axial and lateral interactions within homologous filaments dictate the signaling specificity and order of the AIM2-ASC inflammasome. Nat Commun 12, 2735 (2021).

9. Shen, C. et al. Phase separation drives RNA virus-induced activation of the NLRP6 inflammasome. Cell 184, 5759–5774 e20 (2021).

10. Roberts, T.L. et al. HIN-200 proteins regulate caspase activation in response to foreign cytoplasmic DNA. Science 323, 1057–60 (2009).

11. Barnett, K.C., Li, S., Liang, K. & Ting, J.P. A 360 degrees view of the inflammasome: Mechanisms of activation, cell death, and diseases. Cell 186, 2288–2312 (2023).

12. Jin, T. et al. Structures of the HIN domain:DNA complexes reveal ligand binding and activation mechanisms of the AIM2 inflammasome and IFI16 receptor. Immunity 36, 561–71 (2012).

13. Ni, X. et al. New insights into the structural basis of DNA recognition by HINa and HINb domains of IFI16. J Mol Cell Biol 8, 51–61 (2016).

14. Unterholzner, L. et al. IFI16 is an innate immune sensor for intracellular DNA. Nat Immunol 11, 997–1004 (2010).

15. Antiochos, B., Matyszewski, M., Sohn, J., Casciola-Rosen, L. & Rosen, A. IFI16 filament formation in salivary epithelial cells shapes the anti-IFI16 immune response in Sjogren’s syndrome. JCI Insight 3(2018).

16. Stratmann, S.A., Morrone, S.R., van Oijen, A.M. & Sohn, J. The innate immune sensor IFI16 recognizes foreign DNA in the nucleus by scanning along the duplex. Elife 4, e11721 (2015).

17. Morrone, S.R. et al. Cooperative assembly of IFI16 filaments on dsDNA provides insights into host defense strategy. Proc Natl Acad Sci U S A 111, E62–71 (2014).

18. Garg, A., Stallings, C.M. & Sohn, J. Filament assembly underpins the double-stranded DNA specificity of AIM2-like receptors. Nucleic Acids Res 51, 2574–2585 (2023).

19. Kerur, N. et al. IFI16 acts as a nuclear pathogen sensor to induce the inflammasome in response to Kaposi Sarcoma-associated herpesvirus infection. Cell Host Microbe 9, 363–75 (2011).

20. Li, T., Diner, B.A., Chen, J. & Cristea, I.M. Acetylation modulates cellular distribution and DNA sensing ability of interferon-inducible protein IFI16. Proc Natl Acad Sci U S A 109, 10558–63 (2012).

21. Orzalli, M.H. et al. cGAS-mediated stabilization of IFI16 promotes innate signaling during herpes simplex virus infection. Proc Natl Acad Sci U S A 112, E1773–81 (2015).

22. Gewaid, H. & Bowie, A.G. The ever-expanding role of IFI16 in the anti-viral innate immune response. Immunol Lett 279, 107139 (2026).

23. Deng, Y. et al. IFI-16 inhibition attenuates myocardial remodeling following myocardial infarction. iScience 27, 110568 (2024).

24. Li, T., Chen, J. & Cristea, I.M. Human cytomegalovirus tegument protein pUL83 inhibits IFI16-mediated DNA sensing for immune evasion. Cell Host Microbe 14, 591–9 (2013).

25. Orzalli, M.H., Conwell, S.E., Berrios, C., DeCaprio, J.A. & Knipe, D.M. Nuclear interferon-inducible protein 16 promotes silencing of herpesviral and transfected DNA. Proc Natl Acad Sci U S A 110, E4492–501 (2013).

26. Lee, H.R. et al. Kaposi’s sarcoma-associated herpesvirus viral interferon regulatory factor 4 (vIRF4) targets expression of cellular IRF4 and the Myc gene to facilitate lytic replication. J Virol 88, 2183–94 (2014).

27. Almine, J.F. et al. IFI16 and cGAS cooperate in the activation of STING during DNA sensing in human keratinocytes. Nat Commun 8, 14392 (2017).

28. Chang, X. et al. The role of IFI16 in regulating PANoptosis and implication in heart diseases. Cell Death Discov 10, 204 (2024).

29. Justice, J.L. et al. Systematic profiling of protein complex dynamics reveals DNA-PK phosphorylation of IFI16 en route to herpesvirus immunity. Sci Adv 7(2021).

30. Jonsson, K.L. et al. IFI16 is required for DNA sensing in human macrophages by promoting production and function of cGAMP. Nat Commun 8, 14391 (2017).

31. Jakobsen, M.R. & Paludan, S.R. IFI16: At the interphase between innate DNA sensing and genome regulation. Cytokine Growth Factor Rev 25, 649–55 (2014).

32. Lo Cigno, I., et al. The Nuclear DNA Sensor IFI16 Acts as a Restriction Factor for Human Papillomavirus Replication through Epigenetic Modifications of the Viral Promoters. J Virol 89, 7506–20 (2015).

33. Hotter, D. et al. IFI16 Targets the Transcription Factor Sp1 to Suppress HIV-1 Transcription and Latency Reactivation. Cell Host Microbe 25, 858–872 e13 (2019).

34. Jiang, Z. et al. IFI16 directly senses viral RNA and enhances RIG-I transcription and activation to restrict influenza virus infection. Nat Microbiol 6, 932–945 (2021).

35. Jiang, H., Swacha, P. & Gekara, N.O. Nuclear AIM2-Like Receptors Drive Genotoxic Tissue Injury by Inhibiting DNA Repair. Adv Sci (Weinh) 8, e2102534 (2021).

36. Dunphy, G. et al. Non-canonical Activation of the DNA Sensing Adaptor STING by ATM and IFI16 Mediates NF-kappaB Signaling after Nuclear DNA Damage. Mol Cell 71, 745–760 e5 (2018).

37. Gamble, A. et al. IFI16 senses and protects stalled replication forks. Mol Cell 86, 258–272 e6 (2026).

38. Antiochos, B. et al. The DNA sensors AIM2 and IFI16 are SLE autoantigens that bind neutrophil extracellular traps. Elife 11(2022).

39. Liu, D., Yang, J. & Cristea, I.M. Liquid-liquid phase separation in innate immunity. Trends Immunol 45, 454–469 (2024).

40. Liu, D. et al. IFI16 phase separation via multi-phosphorylation drives innate immune signaling. Nucleic Acids Res 51, 6819–6840 (2023).

41. Huang, X., Zhang, Y., Zhao, Z. & Li, X. Unified phase separation drives DNA-induced inflammasome activation of the AIM2-like receptors. Cell Rep 45, 117065 (2026).

42. Halma, M.T.J. & Xu, L. Life under tension: the relevance of force on biological polymers. Biophys Rep 10, 48–56 (2024).

43. Bustamante, C., Bryant, Z. & Smith, S.B. Ten years of tension: single-molecule DNA mechanics. Nature 421, 423–7 (2003).

44. Alberti, S. et al. Current practices in the study of biomolecular condensates: a community comment. Nat Commun 16, 7730 (2025).

45. Chin Sang, C., Upadhyay, S., Nosella, M.L., Forman-Kay, J.D. & Lee, H.O. The dynamic and heterogeneous composition of biomolecular condensates and its functional relevance. Nat Rev Mol Cell Biol 27, 278–296 (2026).

46. Pappu, R.V., Cohen, S.R., Dar, F., Farag, M. & Kar, M. Phase Transitions of Associative Biomacromolecules. Chem Rev 123, 8945–8987 (2023).

47. Geertsema, H.J. et al. Single-molecule imaging at high fluorophore concentrations by local activation of dye. Biophys J 108, 949–56 (2015).

48. Garg, A. et al. Structural insights into the atypical filament assembly of pyrin domain-containing IFI16. EMBO J 44, 7702–7720 (2025).

49. Wang, P.H. et al. Inhibition of AIM2 inflammasome activation by a novel transcript isoform of IFI16. EMBO Rep 19(2018).

50. Wu, S. et al. Design principles for inflammasome inhibition by pyrin-only-proteins. Elife 13(2024).

51. Gien, H. et al. Cationic Residues of the HIV-1 Nucleocapsid Protein Enable DNA Condensation to Maintain Viral Core Particle Stability during Reverse Transcription. Viruses 16(2024).

52. Ahlawat, V., Mudiyanselage, H.E., Kota, D. & Zhou, H.X. Measuring bridging forces in protein-DNA condensates. Proc Natl Acad Sci U S A 123, e2523058123 (2026).

53. Chappidi, N. et al. PARP1-DNA co-condensation drives DNA repair site assembly to prevent disjunction of broken DNA ends. Cell 187, 945–961 e18 (2024).

54. Syrjanen, J.L. et al. Single-molecule observation of DNA compaction by meiotic protein SYCP3. Elife 6(2017).

55. Gutierrez-Escribano, P. et al. A conserved ATP-and Scc2/4-dependent activity for cohesin in tethering DNA molecules. Sci Adv 5, eaay6804 (2019).

56. Wang, M.D., Yin, H., Landick, R., Gelles, J. & Block, S.M. Stretching DNA with optical tweezers. Biophys J 72, 1335–46 (1997).

57. Ka, N.L., Lim, G.Y., Hwang, S., Kim, S.S. & Lee, M.O. IFI16 inhibits DNA repair that potentiates type-I interferon-induced antitumor effects in triple negative breast cancer. Cell Rep 37, 110138 (2021).

58. Johnstone, R.W., Wei, W., Greenway, A. & Trapani, J.A. Functional interaction between p53 and the interferon-inducible nucleoprotein IFI 16. Oncogene 19, 6033–42 (2000).

59. Shen, Z. et al. Biological condensates form percolated networks with molecular motion properties distinctly different from dilute solutions. Elife 12(2023).

60. Wang, Z. & Zhang, H. Phase Separation, Transition, and Autophagic Degradation of Proteins in Development and Pathogenesis. Trends Cell Biol 29, 417–427 (2019).

61. Lueck, J. et al. Dimerization-dependent gel-like condensation with dsDNA underpins the activation of human cGAS. Cell Rep 45, 116920 (2026).

62. Jiang, Z. et al. Crucial role of the cGAS N terminus in mediating flowable and functional cGAS-DNA condensate formation via DNA interactions. Proc Natl Acad Sci U S A 122, e2411659122 (2025).

63. Zhang, Y. et al. DNA translocation and loop formation mechanism of chromatin remodeling by SWI/SNF and RSC. Mol Cell 24, 559–68 (2006).

64. Zhang, X., Chen, H., Fu, H., Doyle, P.S. & Yan, J. Two distinct overstretched DNA structures revealed by single-molecule thermodynamics measurements. Proc Natl Acad Sci U S A 109, 8103–8 (2012).

65. Wang, H.Y., Elston, T., Mogilner, A. & Oster, G. Force generation in RNA polymerase. Biophys J 74, 1186–202 (1998).

66. Diaz-Celis, C. et al. Assignment of structural transitions during mechanical unwrapping of nucleosomes and their disassembly products. Proc Natl Acad Sci U S A 119, e2206513119 (2022).

67. Mazanek, Z. & Sohn, J. Tracking the polymerization of DNA sensors and inflammasomes using FRET. Methods Enzymol 625, 87–94 (2019).

